# Herb-CMap: Novel Hot Diffusion Algorithm to Identify Bioactive Ingredients from Herbal Medicine Based on Gene Perturbation Profiling

**DOI:** 10.1101/2023.10.19.563046

**Authors:** Yinyin Wang, Yihang Sui, Jiaqi Yao, Jiang Hong, Qimeng Tian, Yun Tang, Jing Tang, Ninghua Tan

## Abstract

Herbal medicine, especially Traditional Chinese medicine (TCM), is a valuable resource of natural products for drug discovery with obvious therapeutic effects. However, the mechanisms of action (MOAs) of herbal medicine are often unknown due to limited target information and the complexity of multiple ingredients and targets. This study developed a Herb-CMap algorithm to prioritize active ingredients and targets within herbal medicine by integrating transcriptomics-based gene perturbation data with a random walk algorithm. This methodology bridges the gap between gene perturbation of herbal medicine and its therapeutic target. Using the Suhuang antitussive capsule (Suhuang) for treating cough variant asthma (CVA) as a case study, we identify and experimentally verify that quercetin and luteolin directly interact with Il17a, Pik3cb, Pik3cd, Akt1, and Tnf. These interactions inhibit the IL-17 signaling pathway and inactivate PI3K, Akt, and NF- κB, preventing lung inflammation and treating CVA. Our findings demonstrate the potential of the Herb-CMap methodology to provide insights into the molecular MOAs of herbal medicine, thereby advancing drug discovery from herbal medicine.

## 1. Introduction

Traditional Chinese medicine (TCM) has been extensively used in China and other countries, presenting exclusive advantages in drug discovery as significant resources. TCM prescriptions adopt the strategy of multiple ingredients, targets, and pathways to achieve optimal therapeutic effects ^1^. However, besides complex ingredients, there could also be many drug-drug interactions in one TCM prescription, making it challenging to clarify the molecular MOAs of TCM prescriptions *via* wet experiments or human experience.

With the development of computational methods, various cutting-edge techniques have been applied to discover the MOAs of herbal medicine, such as molecular docking, machine learning methods ^2,3^, and network-based methods ^4,5^. In particular, network-based methods can extract internal relations according to network topological characteristics, thus revealing hidden relationships, such as drug-disease associations and drug-drug interactions ^6^. The premise behind network-based methods is the observation that genes with similar phenotypes and related functions are located close to each other in the protein-protein interaction (PPI) network. However, in these traditional strategies for network analysis in herbal medicine, the interactions between two ingredients and two herbs are usually ignored ^7^. Recently, Cheng et al. proposed a novel methodology for analyzing drug combinations and drug-disease associations by calculating the overall network distance between the subnetwork of their targets in PPI ^8^. We applied the network distance model to identify herb pairs and quantify their interactions and showed that the network distance algorithm can better capture the biological interactions between herb ingredients ^9^.

As we know, one drug could create effects by binding to certain therapeutical targets and thus causing downstream cascade reactions. Despite the advantages of network methods in herbal medicine, the lack of reliable target information on ingredients is one of the most challenging problems^10^. Due to the development of high-throughput sequencing and system biology, multi-omics data are often integrated into computational target predicting methods, such as molecular docking ^11^, machine learning methods ^12^, and network-based methods ^13^, to improve prediction accuracy ^14^, especially the network perturbation method. For example, the random walk algorithm can be used to prioritize the critical drug targets of the compound based on the topological characteristics of the horizontal disturbance of the gene transcriptome among the PPI network ^15^. Xiao et al., for instance, used differentially expressed genes to establish the weighted gene co-expression network to obtain the hub genes^16^. Another network perturbation called local radiality was developed to prioritize the targets by integrating topological data and the perturbed gene information ^17^. However, as the application domain of these methods was designed to determine drug-drug or drug-disease associations, they have rarely been applied to analyze the complex systems of herbal medicine. Nevertheless, multi-omics integration methods have achieved considerable success in the study of herbal medicine^18^, for instance, interpreting TCM syndrome classifications by integrating proteomic and metabolomic data^19^, exploring the therapeutic effects of bioactivity ingredients in herbs by integrating lipidomic and transcriptomic data^20^, and identifying diagnostic markers of herbal medicine based on network pharmacological analysis of TCM metabolomic data^21^. These exciting findings show the potential of integrating multi-omics data to explore the MOAs of herbal medicine.

Cough variant asthma (CVA) is a type of asthma with bronchial hyperreactive eosinophilic airway inflammation and airway remodeling. Severe and frequent coughs will damage the respiratory tract and aggravate respiratory infection, which has affected patients’ quality of life and caused a considerable burden to society ^22,23^. CVA has the features of paroxysmal, spasmodic, and repetitive coughs, which corresponds to the “wind cough” syndrome in TCM theory. The Suhuang antitussive capsule (Suhuang) is one of the TCM prescriptions used to treat the “wind cough” syndrome. Suhuang, which consists of nine herbs, *Ephedrae Herba* (MAHUANG), *Perillae Folium* (ZISUYE), *Pheretima* (DILONG), *Cicadae Periostracum* (CHANTUI), *Arctii Fructus* (NIUBANGZI), *Schisandrae Chinensis Fructus* (WUWEIZI), *Peucedani Radix* (QIANHU), *Eriobotryae Folium* (PIPAYE), and *Perillae Fructus* (ZISUZI), is mainly used to treat cough, throat itching, and other diseases caused by “wind evil invading lung”, especially for CVA and post-infectious cough (PIC). Although Suhuang has significant clinical therapeutic effects, its mechanisms of action remain unclear. Our laboratory has made progress in previous studies on the MOAs of Suhuang for CVA ^19,24–26^. However, despite this progress, the main active ingredients, their specific therapeutic targets, and the MOAs have still not been systematically clarified. Therefore, more comprehensive research is needed to explain the efficacy of Suhuang for treating CVA.

In this study, we have introduced a novel network perturbation method known as Herb-CMap. This approach is designed to effectively identify therapeutic targets and bioactive ingredients within Traditional Chinese Medicine (TCM) prescriptions. Herb-CMap achieves this by integrating transcriptional gene perturbation data with Protein-Protein Interaction (PPI) networks. The schematic representation of our study can be observed in Fig. 1 and Fig. S1, with the following key steps: Initially, we extracted three crucial types of network graphs from publicly available databases, including the TCM-ingredient-target, disease-gene associations, and PPI networks. Subsequently, we employed a random walk algorithm within the PPI network to enhance transcriptional changes. This process assigned scores to each target. A higher perturbation score indicates a significant proximity of the target to the perturbed genes. We also further conducted molecular docking simulations and wet experiments to validate our prioritized targets and their corresponding ingredients. We firmly believe that the Herb-CMap methodology introduced in our study will provide invaluable insights into unraveling the MOAs underlying complex herbal medicine by connecting the gene perturbation of herbal medicine to its key therapeutical targets.

**Fig. 1.**
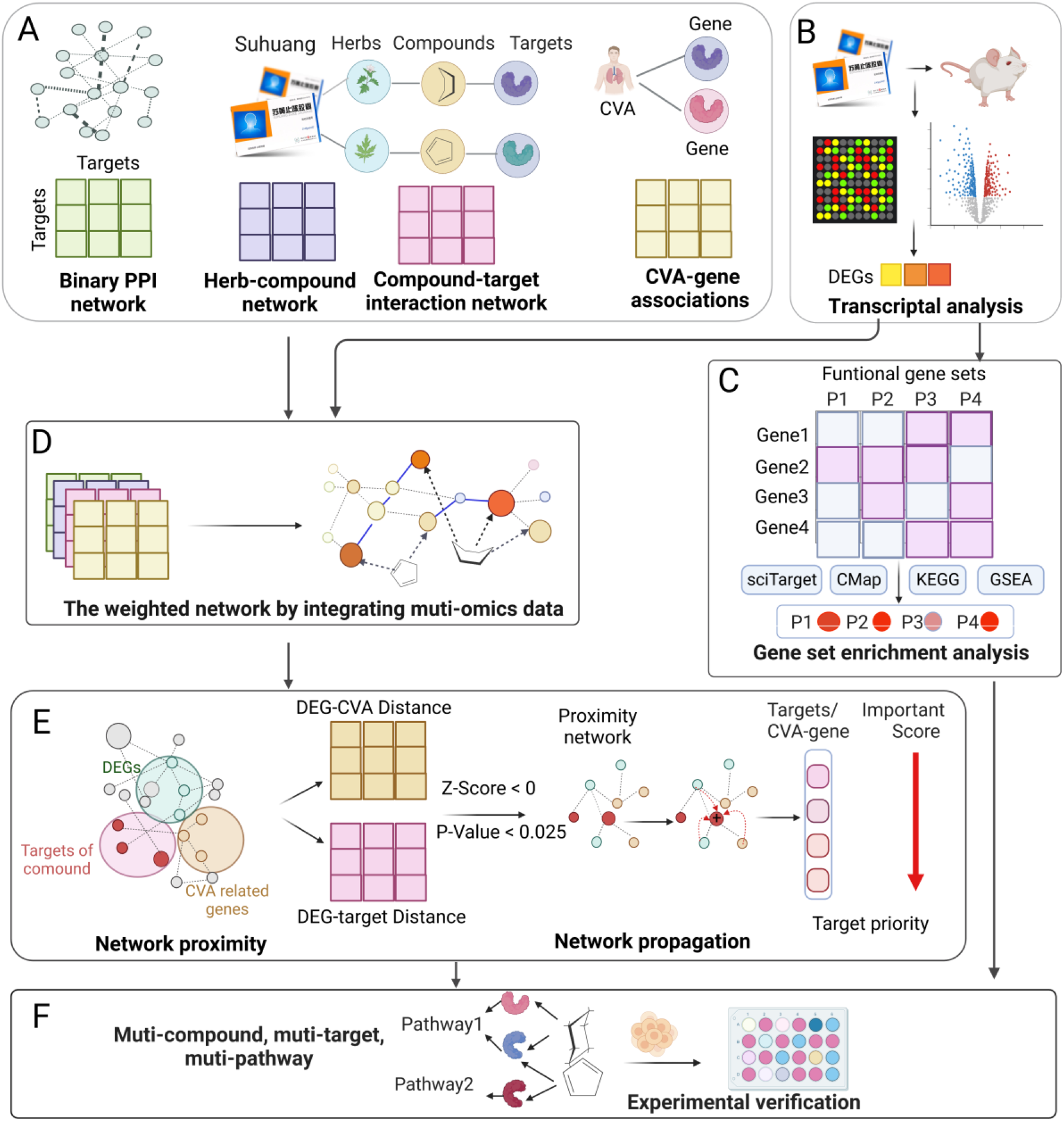
The schema of the Herb-CMap methodology’s sequential steps for connecting gene perturbation to therapeutic targets in the complex system of TCM prescriptions. (A) Data Collection: TCM-ingredient-target relationships, disease-gene associations, and protein-protein interactions (PPIs) were extracted. (B) Gene Perturbation Data: Gene perturbation data were obtained through RNA-seq analysis on CVA rats, identifying differentially expressed genes (DEGs). (C) Functional Enrichment: Various pathway enrichment methods, including KEGG, GSEA, RcisTarget, and CMap, were employed to enrich DEGs with relevant functions at different levels. (D) DEG- Weighted PPI: Gene perturbation data were mapped onto the PPI network to construct a DEG-weighted Protein-Protein Interaction Network. (E) Target-DEG Association Network and network propagation: The proximity between targets and DEGs was calculated to identify vital target-DEG interactions. Network random walk algorithms (LR and hot diffusion) were used to propagate gene perturbation and score each target, with higher gene perturbation scores indicating more essential roles of therapeutic targets. (F) Validation: Prioritized targets and corresponding ingredients were further validated through molecular docking and wet experiments.

In this study, we have introduced an innovative network perturbation method, Herb-CMap. This method is specifically crafted to adeptly identify therapeutic targets and bioactive ingredients within herbal medicine. Herb-CMap accomplishes this task by seamlessly integrating transcriptional gene perturbation data with Protein-Protein Interaction (PPI) networks. The schematic overview of our study is presented in Fig. 1 and Fig. S1, illustrating the following pivotal steps: Initially, we extracted three essential types of network graphs from publicly accessible databases, encompassing the herb-ingredient-target network, disease-gene associations network, and PPI networks; subsequently, we applied a random walk algorithm within the PPI network to amplify transcriptional changes. This procedure assigned scores to each target, with a higher perturbation score signifying a more significant proximity of the target to the perturbed genes; to further validate our prioritized targets and their corresponding ingredients, we use Suhuang for CVA as a case study to identify key target and active ingredients. These findings were verified by both molecular docking simulations and wet experiments, suggesting this approach effectively connects the gene perturbation of herbal medicine to their crucial therapeutic targets, thereby enhancing our understanding of their therapeutic efficacy. We strongly believe that the Herb- CMap methodology introduced in our study will provide invaluable insights into unraveling the Mechanisms of Action (MOAs) that underlie complex herbal medicine.

## 2. Materials and methods

### 2.1 Data collection

The relevant ingredients were collected from six TCM databases, including the Encyclopedia of Traditional Chinese Medicine (ETCM) ^27^, the Traditional Chinese Medicine Systems Pharmacology (TCMSP) ^28^, the Traditional Chinese Medicine Integrated Database (TCMID) ^29^, the Traditional Chinese Medicine on Immuno- Oncology (TCMIO) database ^30^, the Traditional Chinese Medicine Information Database (TCM-ID) ^31^ and Medical Subject Headings (MESH). Only compounds that could be found in the PubChem database ^32^ were kept for further study. Ingredient-target relationships from the STITCH database ^33^ and combined scores > 700 were kept for further study. Subsequently, the properties of components were calculated by ADME-lab ^34^. Disease-related vital targets were also collected. For example, CVA, or “cough variant asthma,” was utilized as a keyword to collect CVA-related genes from the literature. Unlike associated genes from GeneCards ^35^, OMIM ^36^, and DisGeNET ^37^, only those reported as target genes were kept for further study.

The RNA-seq of a TCM prescription can be collected from GEO or manually prepared by experiments. The differentially expressed genes (DEGs) analysis was performed by Deseq2 package in R for further network perturbation calculation.

### 2.2 DEG-weighted PPI network construction

Protein-protein interactions (PPIs) were extracted from the STRING database ^38^. The obtained targets were standardized to Entreze ID by the MyGene package in Python.

First, the binary network was constructed by PPIs with matrix A_N*N_ with N genes; A_i, j_ is the connection between gene *i* and gene *j* with A*_i, j_* _=_ 1 if *i* and *j* are connected else 0.

We obtained the log2 foldchange value of all the genes as a vector F = (f_1_, f_2_, …f_N_) with f_i_ as the log2 foldchange value of a specific gene *i*. The DEG-weighted PPI network (as matrix B) can be established by taking the |log2 foldchange| value of genes as the weight of nodes and the average weight of two nodes as the weight of edges.

### 2.3 Proximity-weighted PPI network construction

Functionally closely related targets tend to be closer to each other in the PPI network, and the shortest path in the network can be used to quantify the interactions of two targets in the DEG-weighted PPI network.

A proximity-weighted PPI network was constructed to quantify the interactions between targets of herb, disease genes, and DEGs. The proximity network for DEGs and disease genes can be defined as matrix C_M×K_. M is the set of DEGs, and K is the set of the targets of TCM prescriptions. We define the distance between any node DEG i in M and node j in K as C_ij_:

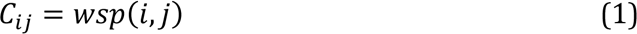

where wsp is the Shortest_path function and it is calculated by the NetworkX package in Python. The pairwise weighted shortest path is calculated as a proximity-weighted PPI network.

Next, we construct our DEG-target association network from matrix C_M×K_ via Z score transformation according to the distance value. Z-scored C_ij_ is calculated for each value in C, and the *p-*value is expressed as Z:

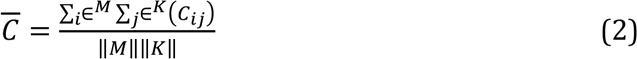

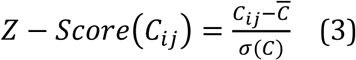

Only target-target pairs with *p*-value < 0.025 were kept connected, and the final transformed proximity-weighted PPI network is defined as D.

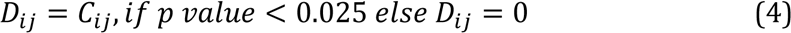

### 2.4 Prioritization of the CVA treatment genes using the local radiality network perturbation method

The union potential targets from multiple ingredients were collected in the TCM prescriptions study. The hypothesis was that the critical targets of active ingredients would activate downstream pathways in a cascade, thus regulating the neighbor targets in the PPI network. On the other hand, the therapeutic disease-related genes would also be significantly regulated by the TCM prescription. They would thus be closely related to the DEGs in the PPI network.

Radiality is a well-known centrality measure describing the level of node reachability of a node via different shortest paths of the network. It was acknowledged that deregulated genes were closer to known targets than most proteins in the network. Based on this hypothesis, local radiality (LR) (Isik et al., 2015) was applied to prioritize the critical disease-associated genes by combining perturbed genes and functional interaction network information. The LR score of node *i* in the network D is calculated as follows:

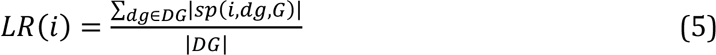

Here, the function |sp| calculates the weighted shortest path that connects the gene dg and the node n in D; |DG| indicates the total number of connected genes. The LR utilizes drug perturbation data (<u>i.e.,</u> connected genes) and topological information (i.e., shortest path distance). Thus, it implements the hypothesis about the proximity of DEGs, targets, and disease genes. In our study, LR (n) represents the critical score of disease genes for causing gene perturbation of TCM prescription/herbal medicine.

### 2.5 Prioritization of the active ingredients and action targets using the random walk method

It was reported that a drug’s targets and DEGs were typically closely connected in PPI within three steps _39_. Here, network perturbation methods are applied to infer the critical targets by aggravating the gene perturbation of all neighborhood DEGs to determine the active ingredients acting on these crucial targets and pathways.

We used a random walk with restart (RWR) algorithm called heat diffusion to screen an essential target query that interacted with DEGs closely. The RWR network perturbation algorithm is described as follows: Diffusion applies heat to each target as the heat source in the target-DEG network, and the heat flows

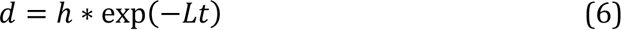

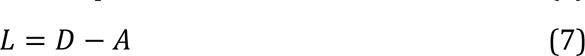

Where “h” is a vector representing the original query or signal. “d” is another vector, which is the result of the diffusion process. The degree matrix D is a diagonal matrix where each diagonal element corresponds to the degree (number of connections) of a node in the graph. The adjacency matrix A encodes the connectivity between nodes. It represents how the original signal has spread or diffused over the network. L is the Laplacian matrix of the network graph and is defined as the difference between the degree matrix (D) and the graph’s adjacency matrix (A). The expression “exp(*)” is the matrix exponential . “t” is a scalar parameter representing the total time of diffusion. It controls how far and quickly the original signal spreads over the network. A larger value of t will allow the signal to spread further, while a smaller value will limit its spread. Here, we conduct a diffusion process on a graph target-DEG network, where “h” represents the transcript gene perturbation values of nodes, and the Laplacian matrix is used to model the diffusion behavior. The parameter ’t’ controls the extent and speed of the diffusion. The result of this process is the vector ’d,’ which represents the gene perturbation score of targets and DEGs after diffusion. We use a default value of t = 0.1 ^40^.

### 2.6 Case study: Identifying the active ingredients and MOAs of Suhuang for CVA

Male Wistar rats were intraperitoneally injected with ovalbumin (OVA) to establish the CVA rat model ^26^. The rats from the experimental group were intragastrically administrated 7 g/kg of Suhuang crude drug for two weeks. Subsequently, the rats were sacrificed by euthanasia. The lungs from the control, CVA, and CVA+Suhuang groups were collected and subjected to transcriptome sequencing.

The Deseq2 package in R. Gene obtained the differentially expressed genes (DEGs) analysis set enrichment analysis (GSEA) based on Go Ontology (GO) and Kyoto Encyclopedia of Genes and Genomes (KEGG) datasets to understand these DEGs’ biological functions and related pathways. RcisTarget transcription factor (TF) analysis was conducted to find the critical genes to determine the key DEGs’ specific regulated mechanisms. Moreover, the Connectivity Map (CMap) was adopted to excavate the functional connections between CVA and the genetic perturbation of Suhuang ^41^. In order to study whether the gene expression of DEGs in different tissues is consistent with the meridian classifications of Suhuang in TCM theory, the gene expression of DEGs at the transcription level was extracted from the Human Protein Atlas databases. More detail about the CVA model and average RNA data analysis for Suhuang can be seen in **Supplementary Note 1**. Finally, the Herb-CMap method was used to prioritize Suhuang’s crucial genes and targets for CVA.

### 2.7 Verifying the critical targets and active ingredients

*Verification of targets and pathways:* RAW264.7 cells were plated in 6-well plates at a density of 2 × 10^6^ cells and pretreated with different concentrations of Suhuang (0.1 μg/ml, 0.2 μg/ml, 0.4 μg/ml), and then stimulated with 1 μg/ml LPS for six h. Total RNA was isolated from cells using Trizol reagent (Takara, Otsu, Japan) according to the manufacturer’s protocol. RNAs were reverse transcribed to cDNAs for quantitative PCR. The primer sequences were as follows:

**Table 1.**
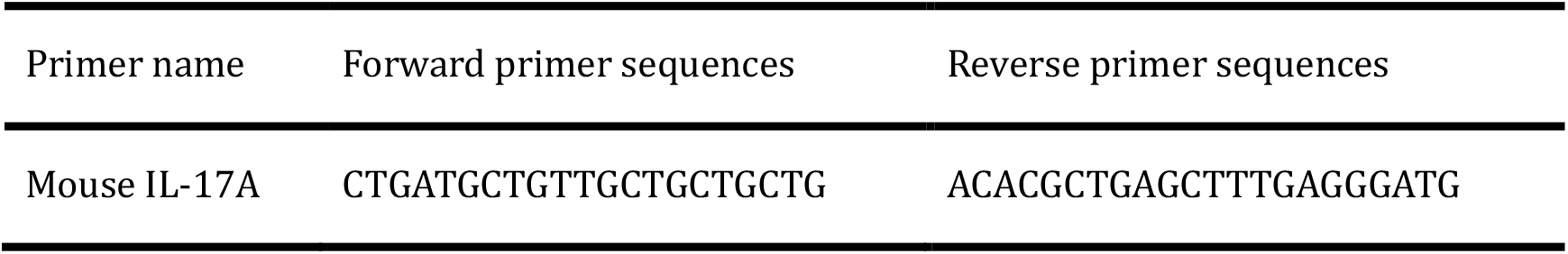
Sequences of the primers used for real-time PCR.

*Validation of active ingredient and target interaction:* We used AutoDock Vina for molecular docking ^42^. According to the scores of MCODE ^43^, the core targets of Suhuang for treating CVA were selected. Then, their crystal structures were downloaded from the RCSB Protein Data Bank (PDB) ^44^. Next, these crystal structures were modified by PyMol to remove the original ligands and water molecules and then saved in the PDBQT format. The three-dimensional structures of related compounds were downloaded from PubChem. Then, the bond energy of the compound ligand molecules was processed via Chem3D 15.0 to reduce intermolecular repulsion and saved in the PDB format. Finally, the three-dimensional crystal structures of target proteins and corresponding compound ligands were introduced into AutoDock Vina.

*Verification of active ingredients:* RAW264.7 cells were plated in 96-well plates at a density of 1×10^4^ cells. RAW264.7 cells were treated with LPS (1 μg/mL) and different concentrations of luteolin, quercetin, and Dexamethasone (2.5, 5, 10, 20, 40, 80 μM). Then, the cells were incubated for 24 h, and the content of NO in cell culture supernatant was determined by using the Griess reaction. The NO inhibition rate of different drugs was calculated according to the following formula:

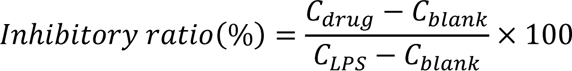

where *C*_blank_ is the content of NO in unprocessed cell culture, *C_LPS_* is the content of NO in LPS-induced cell culture, and *C_drug_* is the content of NO in cell culture treated with LPS and different concentrations of drugs. The IC_50_ values for luteolin, quercetin, and Dexamethasone were determined on three replicates of 6 concentrations. Statistical analysis and the IC_50_ values were calculated using GraphPad Prism version 8.0.2 software from GraphPad Software Incorporated.

## 3. Results

### 3.1 Data statistics on the herb-ingredient-target network of Suhuang

In total, 1,377 ingredients were collected from six TCM databases. The overlapping ingredients among these six databases are shown in Fig. 2A. The number of ingredients varied among different databases, with 849 ingredients from TCMIO, followed by TCMSP, and only 250 ingredients from ETCM. Furthermore, the TCMSP and TCMIO databases shared the most common compounds (n = 406). Primarily, we found only 14 compounds appearing in all six databases, suggesting the necessity of data integration to address the data complexity.

**Fig. 2.**
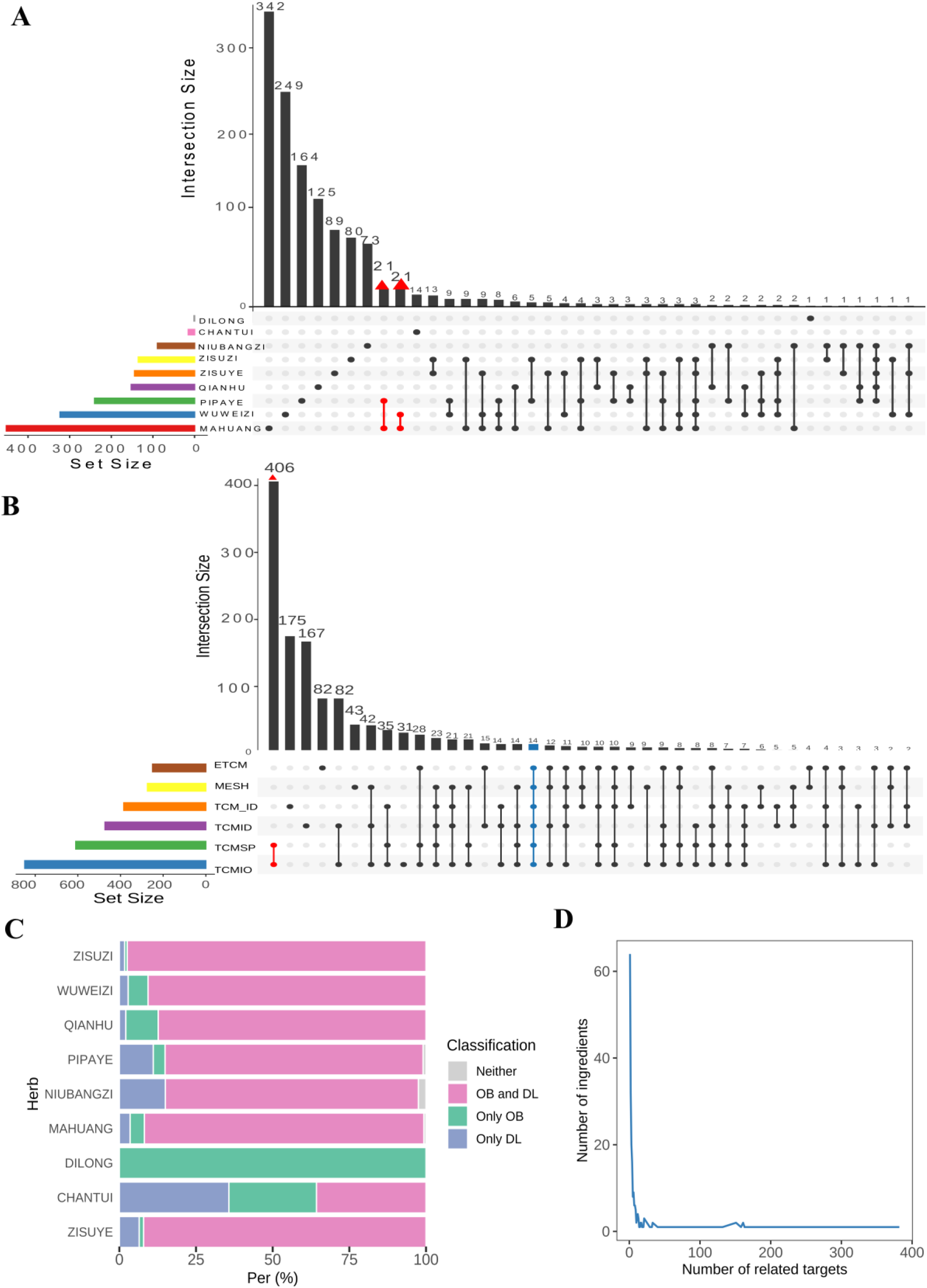
Data Statistics of the Herb-Ingredient-Target Network. (A) The overlapping situation of the ingredients from Suhuang. **(B)** The overlapping of the ingredients from different databases. **(C)** The percentage of OB and DL for ingredients in Suhuang. **(D)** The distribution of the number of targets for each ingredient in Suhuang.

The ingredients in Suhuang are derived from nine herbs, with N*_MAHUANG_* = 452, N*_WUWEIZI_* = 323, N*_PIPAYE_* = 240, N*_QIANHU_* = 152, N*_ZISUYE_* = 144, N*_ZIZUZI_* = 135, N*_NIUBANGZI_* = 89, N*_CHANTUI_* = 15, and N*_DILONG_* = 1(Fig. 2B). MAHUANG shared the most common components with PIPAYE and WUWEIZI (n = 21 and 21 separately). Except for two animal drugs, DILONG and CHANTUI, the other seven herbs have a high percentage of compounds with good OB and DL ranging from around 80% to 95% (Fig. 2C). As shown in Fig. 2D, 60.6% of ingredients were predicted to bind with less than five targets of the compounds. Only 63 ingredients owned more than ten targets.

### 3.2 DEG analysis and their tissue distribution

To grasp the gene perturbation of Suhuang for CVA, RNA-seq data were tested in three groups: control, CVA, and Suhuang. Compared to the control group, a total of 589 genes were identified as differentially expressed genes (DEGs, *p*.adjust < 0.05 and |log2 foldchange| > 2) for the CVA group, with 465 upregulated and 124 downregulated (Fig. 3A). Furthermore, a total of 790 genes were identified as DEGs when comparing the Suhuang group to the CVA group, with 300 upregulated and 490 downregulated (Fig. 3B). In total, 338 common DEGs were identified between CVA/Control and Suhuang/CVA (Fig. 3C), suggesting that these abnormal DEGs in the disease condition returned to the normal level after the treatment of Suhuang (Fig. 3D).

**Fig. 3.**
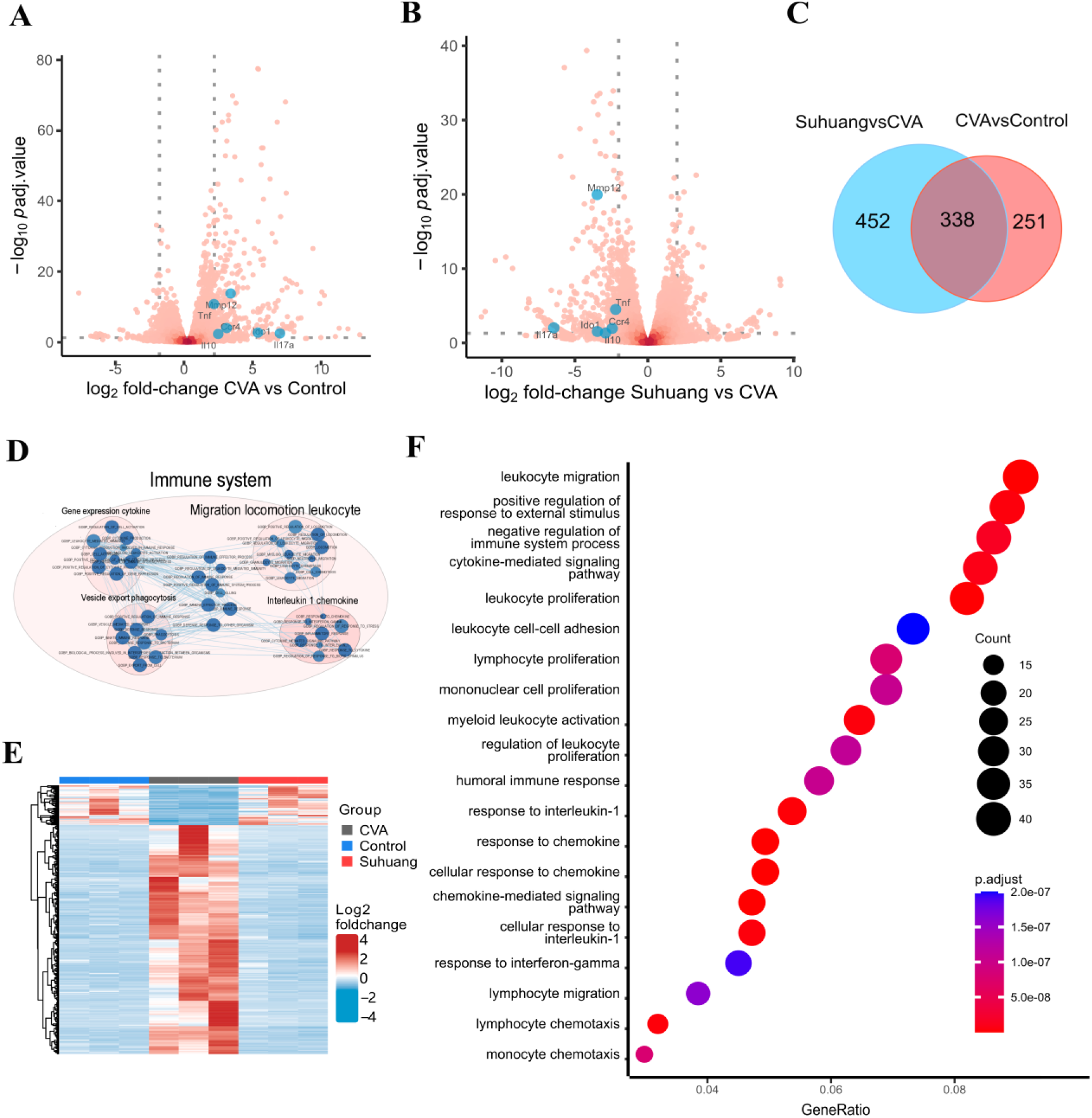
Transcriptomics analysis of Suhuang-treated OVA-induced CVA rats. **(A)** DEGs volcano plot of Suhuang *versus*. CVA. **(B)** DEGs volcano plot of CVA *versus*. Control. **(C)** The Venn diagram of DEGs. **(D)** The hierarchical clustering heat map of DEGs. **(E)** The visual analysis of the pathway enrichment results of GSEA analysis for GO: BP gene sets by the plug-in EnrichmentMap of Cytoscape. **(F)** The GO: BP pathway enrichment of the DEGs of Suhuang *versus*. CVA.

Fig. 4A shows the distribution of TCM properties of the nine herbs in Suhuang, including four natures, five flavors, 12 kinds of meridian classifications, and six main efficacies. Regarding the nature of Suhuang, it has warm and cold medicinal characteristics. Regarding the flavors, pungent was the most frequent. In TCM theory, the pungent flavor tended to affect the surface and upper parts of the human body. Furthermore, CVA is a respiratory disease. Accordingly, all herbs in Suhuang were classified to the lung meridian except DILONG. These herbs, especially MAHUANG and CHANTUI, have been reported to have extensive effects, such as refrigeration, analgesia, relieving wheeling, immunological enhancements, immuno-suppression, and hypoglycemia.

**Fig. 4.**
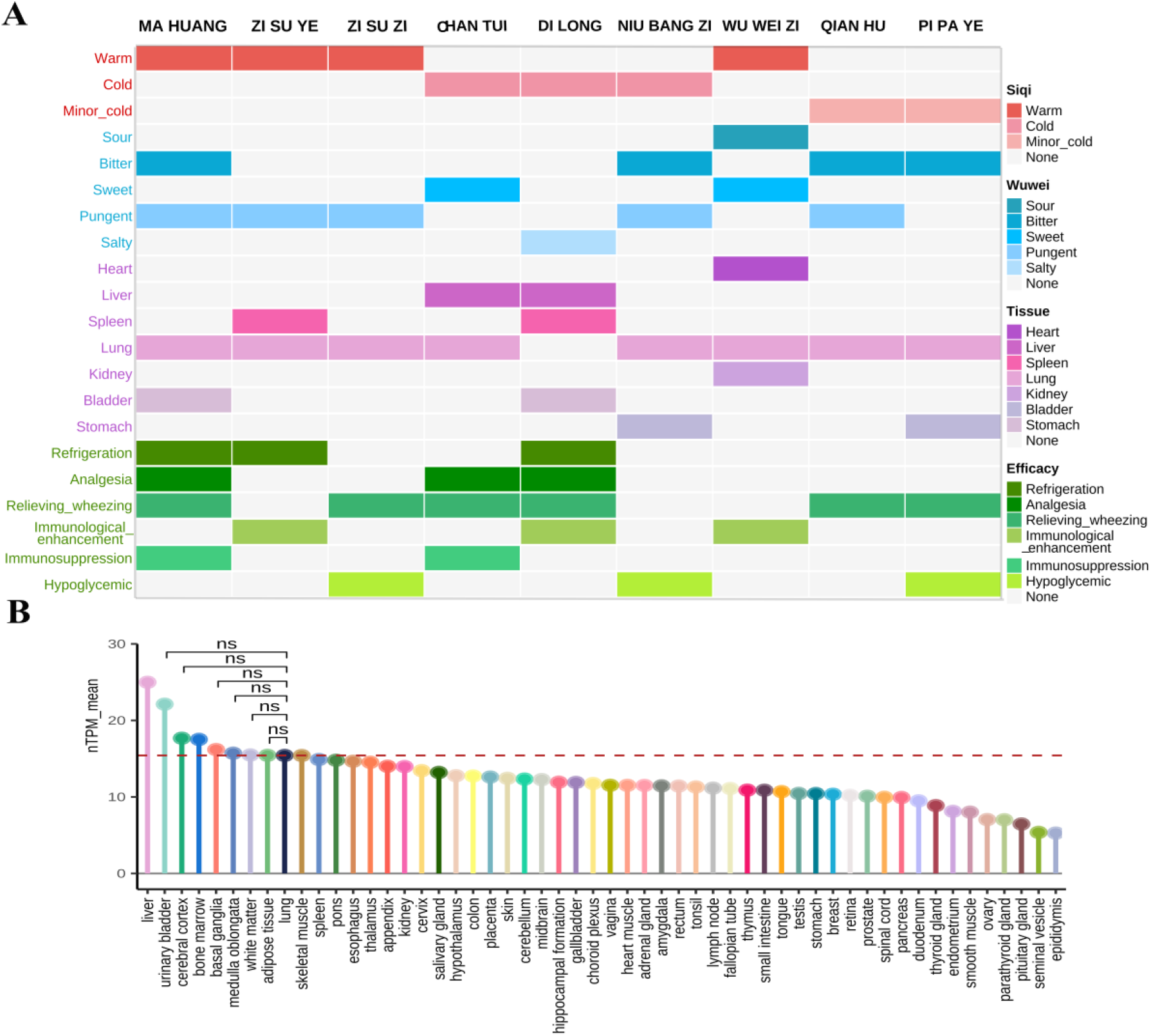
The associations between the tissue-specific expression of DEGs and the meridian types of herbs in Suhuang. (A) The properties distributions of herbs in Suhuang. **(B)** The average expression of DEGs in different tissues.

Considering that the meridian classifications in the TCM theory might provide a reasonable interpretation of Suhuang’s therapeutic effects on CVA, we further investigated whether the tissue distributions of these DEGs at the transcription level were consistent with the meridian distribution of Suhuang from TCM theory. As shown in Fig. 4B, the lung was one of the top 10 highest averaged expressed tissues, and the top 10 tissues were not significantly different. On the other hand, except for the top 10 genes, the expression of DEGs in normal human lung tissue was significantly higher than the other 53 tissues. Interestingly, the expression of the DEGs in the liver and urinary bladder tissues was the highest, which was consistent with the liver meridian of CHANGTUI and DILONG and the bladder meridian of MAHUANG and DILONG (Fig. 4A). Taken together, we verified that the results of gene perturbation of Suhuang were in line with the meridian classifications in TCM theory and that Suhuang for treating CVA was closely related to the functional regulation in the lung tissue.

### 3.3 Identifying CVA-related pathways via pathway enrichment analysis

Three enrichment analysis methods, GO, KEGG, and GESA, were performed to systematically grasp these DEGs’ biological functions. We also utilized the CMap and TF analysis to identify the underlying molecular mechanisms further.

In total, 263 GO-biological processes (GO: BP) terms and 26 KEGG pathways were enriched significantly by GO and KEGG analysis, respectively (*p*.adjust value < 0.05, Table S1-2). We also mapped the DEGs onto GSEA analysis for GO: BP gene sets, and the EnrichmentMap plugin in Cytoscape was utilized to identify the hierarchical relationships of these pathways (Table S3). Consequently, the top enriched GO: BP terms for Suhuang were related to inflammatory cytokines in the immune system, such as response to interleukin-1, response to interferon-gamma, and negative regulation of the immune system process (Fig. 3E-F). Therefore, it appears that Suhuang might relieve CVA by extensively regulating the pathways associated with the immune system. According to KEGG pathway enrichment analysis, half of the top 20 significantly enriched pathways were associated with CVA, such as the Asthma pathway, IL-17 signaling pathway, TNF signaling pathway, and Toll-like receptor signaling pathway (Fig. 5A).

**Fig. 5.**
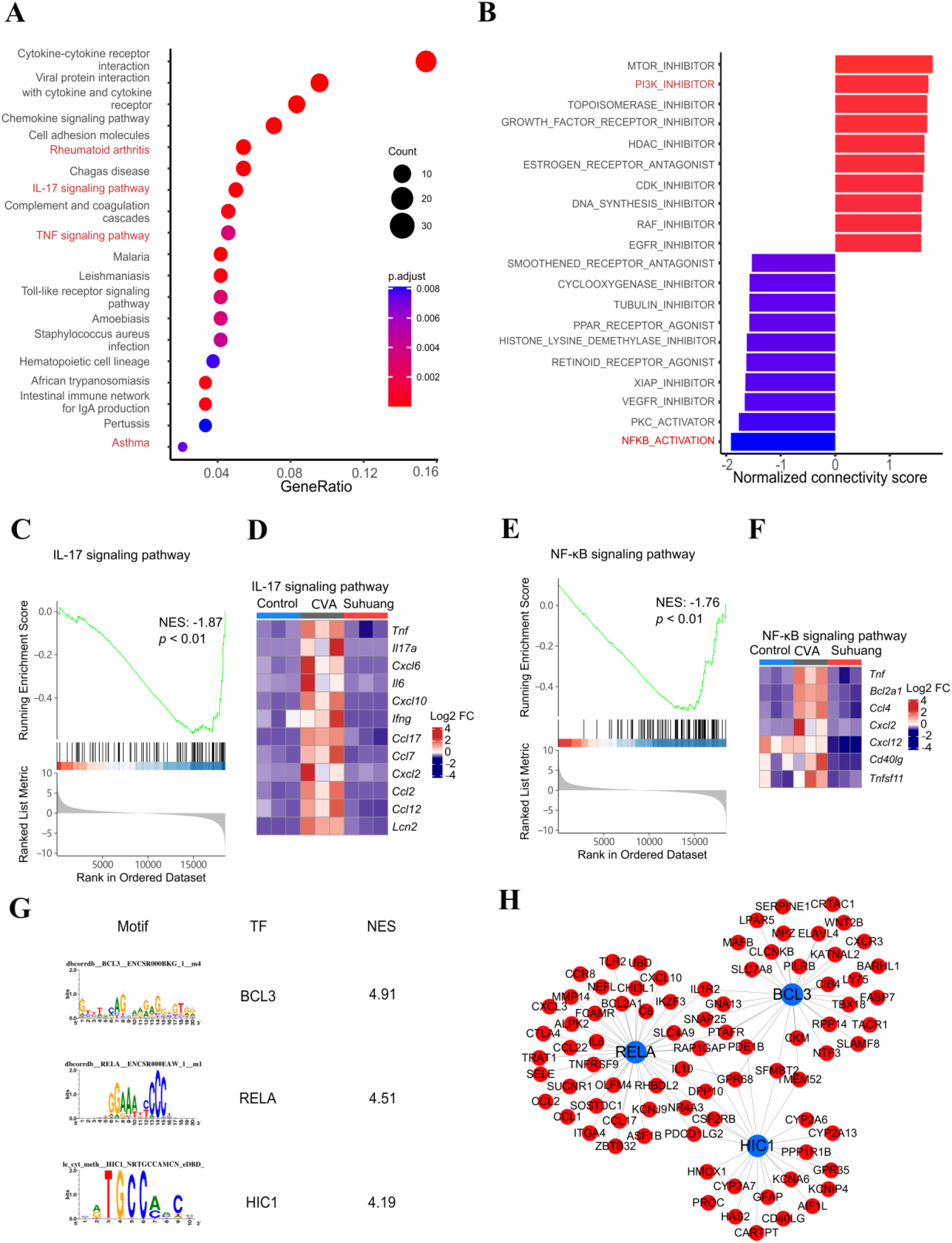
The pathway enrichment analysis of the DEGs in Suhuang *versus*. CVA. **(A)** KEGG pathway enrichment analysis based on the DEGs. **(B)** The gene set enrichment of the Connectivity Map about the modes of action based on the DEGs. The top 10 mechanisms of action (the normalized connectivity score > 0) were positively correlated with Suhuang for treating CVA. **(C)** GSEA plot for the IL-17 Signaling Pathway. **(D)** The expressional heatmap of the DEGs enriched in IL-17 Signaling Pathway. **(E)** GSEA plot for NF-κB Signaling Pathway. **(F)** The expressional heatmap of the DEGs enriched in NF-κB Signaling Pathway. **(G)** The information about the top motif enrichment and its TF annotation. NES: Normalized enrichment score of the enriched motifs in the geneset. The motifs whose NES > 3.0 are considered essential. **(H)** The DEG regulatory network about the motif-TF annotations.

Meanwhile, we mapped the DEGs onto the CMap database to find similar gene signatures. The top and bottom 10 MOAs, respectively, were illustrated in Fig. 5B. The MOAs in Suhuang for treating CVA were positively associated with inhibiting PI3K and negatively associated with activating NF-κB, suggesting that Suhuang may treat CVA through inhibiting PI3K and NF-κB signaling pathway (Fig. 5B, Fig. S2).

Interestingly, both KEGG and GSEA showed that the IL-17 signaling pathway was a significant enrichment pathway in Suhuang for treating CVA (Fig. 5A and C, Table S4). However, it was studied that the IL-17 signaling pathway was closely related to Th1 airway inflammation in asthma, while CVA was generally recognized as Th2 airway inflammation ^45–47^. Therefore, we inferred that Suhuang may have other therapeutic effects on Th1 asthma (Fig. 5D). The NF-κB pathway from GSEA and CMap analysis was also significantly enriched (Fig. 5E and F). In brief, multiple pathway enrichment methods indicated that Suhuang may regulate these pathways to treat CVA in model rats.

According to transcription factor (TF) analysis, 86 motifs were significantly enriched. The result for the top three TFs based on motifs was listed in Fig. 5G. RELA was an essential member of the NF-κB family, with 29 DEGs enriched in this related motif. BCL3, with 31 DEGs enriched in this related motif, was highly similar to the IκB family of NF-κB inhibitors ^48,49^. These two TFs participated in many cellular responses, such as cell proliferation and transformation, apoptosis, inflammation, and immune response, by binding NF-κB or other TFs and regulating target genes (Fig. 5H). These results were consistent with our previous findings about the critical role of the NF-κB signaling pathway in Suhuang for inhibiting non-resolving inflammation and treating acute lung diseases. Our previous work found that Suhuang could relieve acute lung injury by inhibiting the NF-κB signaling pathway ^19^.

### 3.4 Prioritization of the therapeutic genes using the local radiality network perturbation method

Generally speaking, the weighted shortest path (S) between DEGs and CVA genes was negatively correlated with the corresponding |log2foldchange|, suggesting that the more significantly the DEGs expressed, the more functionally close it was to CVA genes (Fig. 6A) in PPI. In addition, DEGs could be classified into three clusters by hierarchical clustering analysis according to the weighted shortest path among 54 disease genes. The weighted shortest path between the CVA genes and DEGs from cluster Ⅱ (average |log2 foldchange|_Ⅱ_=4.086, average S_Ⅱ_ = 1.977) was significantly lower than that of the other two cluster DEGs (average S_Ⅰ, Ⅲ_ = 5.025 and 3.172, average |log2 foldchange| _Ⅰ, Ⅲ_ =2.50,3.10) (Fig. 6B, Fig. 6C), suggesting the importance of the DEGs from cluster Ⅱ . Therefore, further pathway enrichment analysis was performed based on DEGs of cluster Ⅱ. We found that the enriched pathways were similar to those from all the DEGs, suggesting that cluster Ⅱ DEGs could characterize Suhuang perturbation and could be considered as signature genes (Fig. 6D). These results indicated that the weighted proximity network based on DEGs could reveal the vital therapeutic genes of Suhuang for treating CVA.

**Fig. 6.**
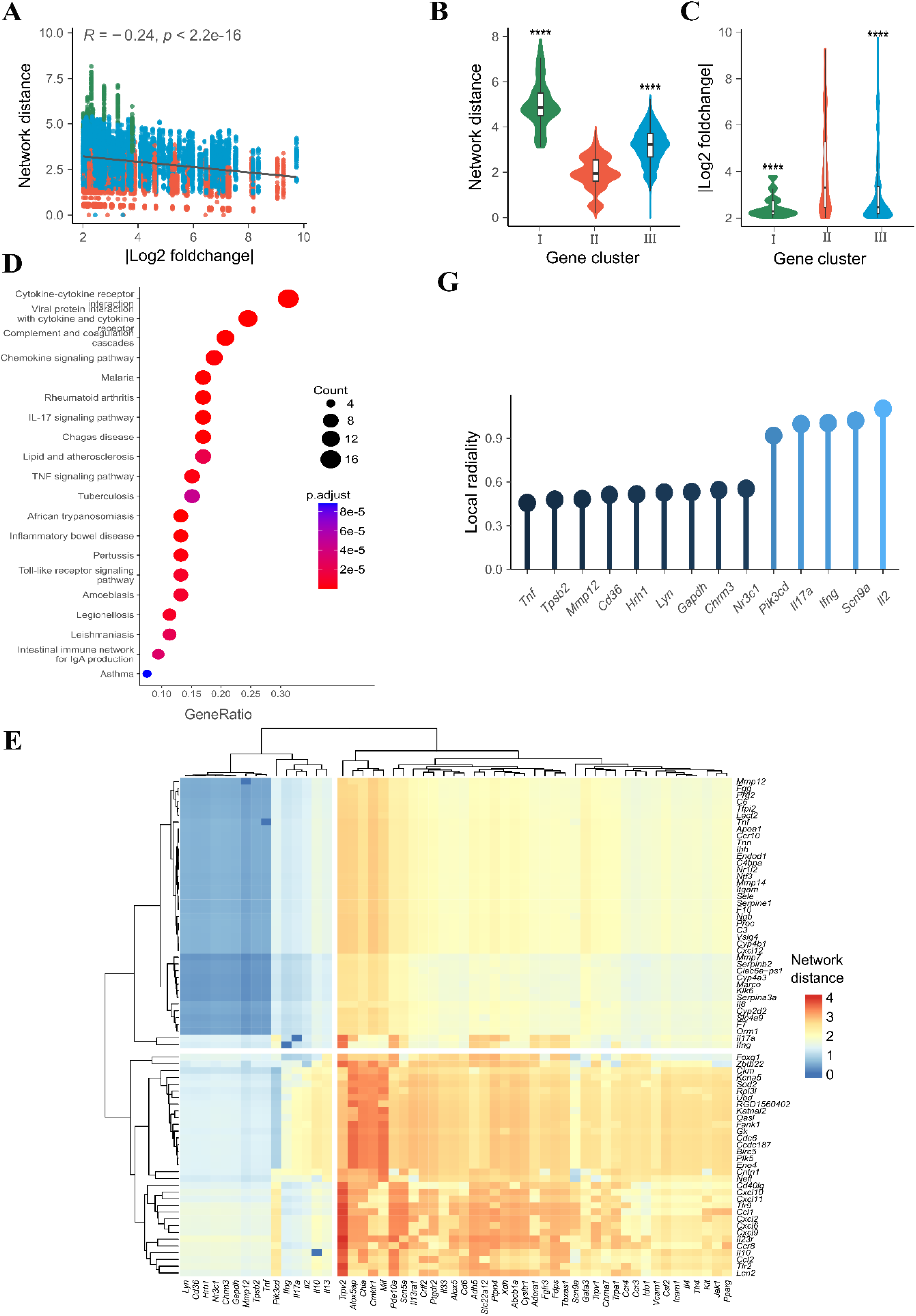
The association between DEGs and CVA-related genes. (A) The scatter plot of the network distance between the DEGs and CVA genes and the log2 foldchange absolute values of the DEGs. The Pearson correlation coefficient (*R*) and two-sided t-distribution *p* values (*p*) for the comparison are shown in the plot. The linear regression trendline (black) and its 95% confidence interval (shaded gray area) are shown in the graph. The colors suggest the clusters of the DEGs. **(B)** Comparison of the network distance between the DEGs of three clusters and CVA genes. *P-values* were calculated by <u>t.test.</u> **\*\*\*\****p* < 0.0001, vs. the cluster Ⅱ group. **(C)** Comparison of the log2 foldchange absolute values of the DEGs of three clusters. **(D)** The top 20 significant signaling pathways were enriched by introducing 73 DEGs into the KEGG pathway enrichment analysis. **(E)** The heatmap of the network distance between 73 DEGs of cluster 2 and CVA genes. The row genes are CVA genes, and the column genes are the DEGs of cluster 2. **(F)** The distribution of the DEGs of cluster 2. The genes of the left nest were significantly enriched in the IL-17 signaling pathway, and the genes of the right nest were enriched in the PI3K-AKt signaling pathway. The genes of the deepest color belonged to CVA genes. **(G)** The LR of the top 14 CVA-related genes with the significantly low network distance (*p*-value < 0.025) is based on the network propagation method.

Then, the LR of network perturbation was leveraged to quantify the importance of disease genes. As shown in Figure 6G, *Tnf* obtained the highest importance score, 0.47, followed by *Tpsb2* (0.48) and *Mmp12* (0.48). *Tnf* was enriched in the IL-17 signaling pathway and was a crucial gene. *Tpsb2* has been implicated as a mediator in the pathogenesis of asthma and other allergic and inflammatory disorders ^50^. *Mmp12*, which encodes a member of the peptidase M10 family of matrix metalloproteinases (MMPs), may play a role in aneurysm formation, and mutations in this gene are associated with lung function and chronic obstructive pulmonary disease (COPD) ^51^.

*Il17a* (score = 1.00) was significantly enriched in the IL-17 signaling pathway, and *Pik3cd* (score = 0.92), enriched in the PI3K-Akt signaling pathway, was also prioritized using our network perturbation method.

Taken together, 15 genes involved in IL-17 and PI3K-Akt signaling pathways were prioritized using network perturbation methods.

### 3.5 Prioritization of the critical targets and corresponding ingredients using the random walk with restart network perturbation method

First, 928,485 target-DEG interactions (T-Gs) were calculated as proximity matrix C_MxN_. Furthermore, 19,009 T-Gs were identified as significant interactions and kept for the proximity network construction (*z* < 0, *p* < 0.025), covering 694 targets (Fig. S4A). We introduced these target genes into the KEGG pathway enrichment analysis. IL-17, PI3K-Akt, and NF-κB signaling pathways were significantly enriched (*p* < 0.01) (Table S1).

With target-DEG interactions, the random walk algorithm was performed to find the key targets. Fig. 7A illustrates the top 20 targets with the highest heat values. Primarily, we found that *Il13*, *Rela* enriched in the IL-17 signaling pathway and *Ywhab*, *Rela*, *Pik3cb* enriched in the PI3K-Akt signaling pathway are among the top 20 targets (Fig. 7A), suggesting Suhuang may influence these two pathways to relieve CVA.

**Fig. 7.**
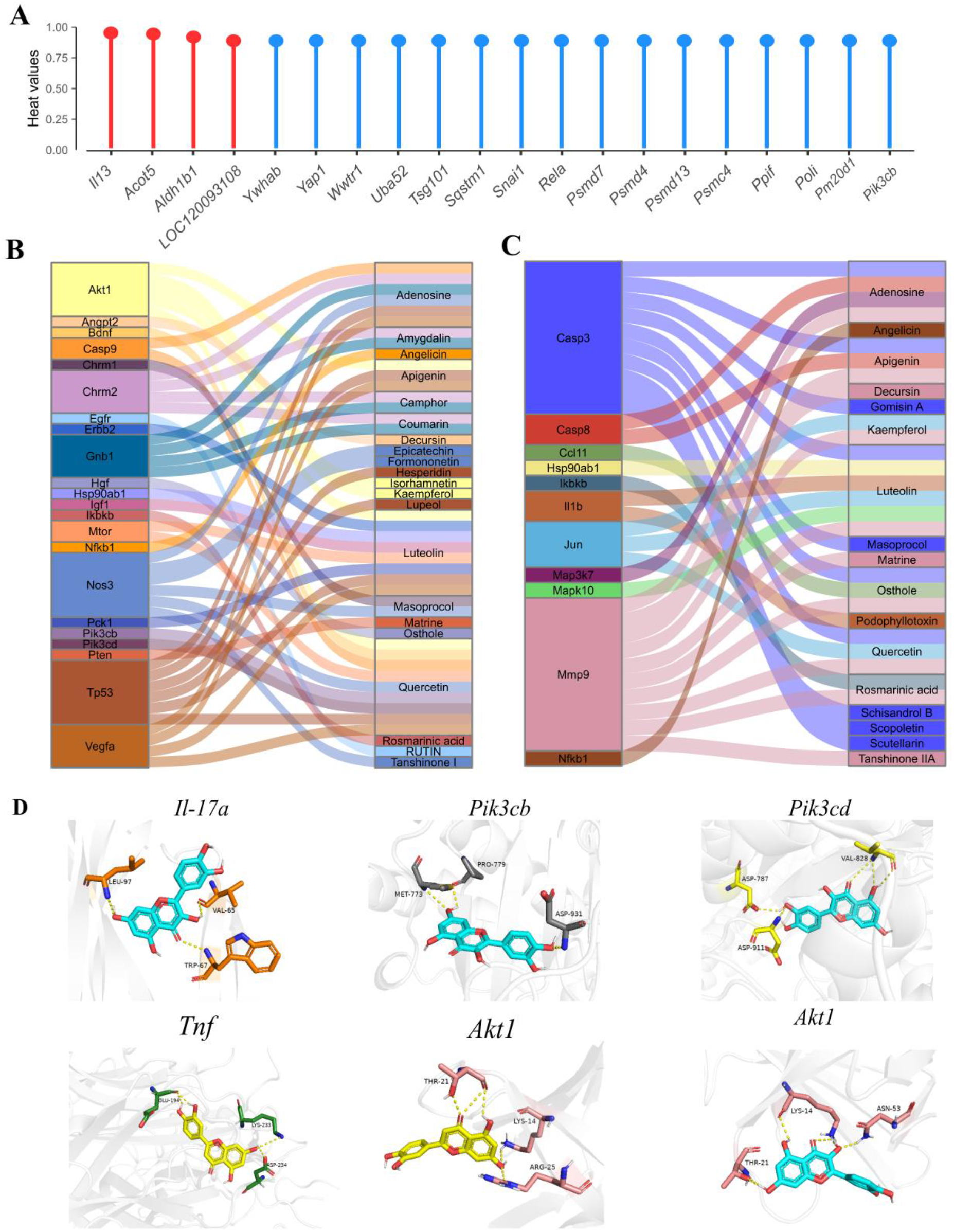
The main active ingredients and the molecular mechanisms of Suhuang for treating CVA. (A) The heat and ranking values of the top 20 targets in the target-DEG interactions by the Cytoscape implementation of Diffusion. The blue genes represented non-differentially expressed genes, and the red genes represented DEGs. (B-C) Sankey diagram of the active ingredients interacted closely with the enriched genes in PI3K-AKt signaling pathway **(B)** and Il-17 signaling pathway **(C)** based on the KEGG pathway enrichment analysis. Active ingredients (right) were connected to their targets (left) according to the compound-target interactions in Suhuang previously mentioned. **(D)** Mocular docking of quercetin on *Il-17a*, *Pik3cb*, *Pik3cd and Akt1* as well as luteolin on *Tnf* and *Akt1*. The essential residues with five colors represented corresponding target genes. Quercetin was shown in blue cartoon structures and luteolin in yellow cartoon structures. The polar interactions between the compounds and the target genes were shown in yellow dashes.

Interestingly, 499 of 694 targets were not significantly changed at the transcription level and were kept in our proximity network. More importantly, according to the heat values from network perturbation, 16 of the top 20 targets were not DEGs, such as *Rela*, *Ywhab*, and *Pik3cb* (Fig. 7A). These results suggested that our network inference methods could reveal potential targets that were not significantly expressed at the transcription level. We also investigated the functions of these 499 targets via KEGG pathway enrichment. We found that the PI3K-Akt and IL-17 signaling pathways were significantly enriched again (Fig. S4B). Forty-five targets were enriched in the PI3K-Akt signaling pathway, and 23 were enriched in the IL-17 signaling pathway.

### 3.6 Experimental validation of the active ingredients and corresponding targets

To identify active ingredients, 23 and 45 targets enriched in IL-17 and PI3K-Akt signaling pathways were selected as critical targets. Correspondingly, molecular docking was performed on the 17 and 21 active compounds predicted to interact with these targets (Fig. 7B-C), respectively. The enriched targets and corresponding ingredients, as well as binding interactions, were shown in Fig. 7D. In particular, quercetin represented low binding energy with *Il17a*, *Pik3cb*, *Pik3cd*, and *Akt1* as -7.4, -7.6, -8.8, -6.9 kcal/mol, respectively. In addition, luteolin also showed stable docking results with *Tnf* and *Akt1,* whose binding energy values were -8.7 and -6.8 kcal/mol, respectively. The above results illustrated that quercetin and luteolin could alleviate lung inflammation by regulating IL-17 and PI3K-Akt signaling pathways and might be the main active compounds of Suhuang for treating CVA.

More importantly, according to Real-time PCR, we found that the mRNA expression of *Il17a* was reduced by Suhuang significantly (Fig. 8A). Quercetin and luteolin showed lower IC_50_ than positive control drug Dexamethasone (Fig. 8B).

**Fig. 8.**
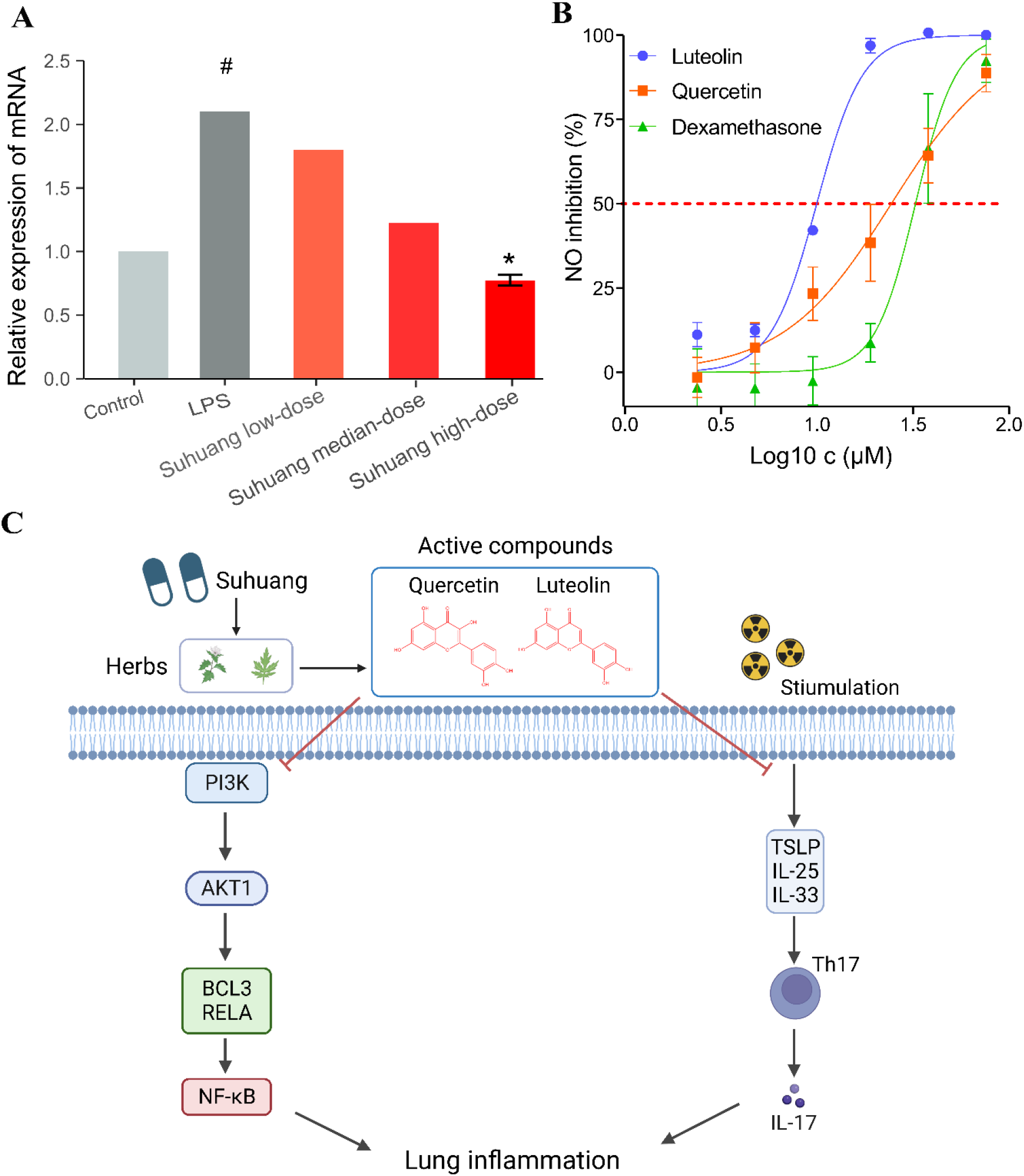
The novel mechanism of Suhuang for CVA treatment was identified by our Herb-CMap. (A) Real-time PCR determined the relative mRNA level of *Il17a* in LPS-induced RAW264.7 cells treated by Suhuang. Data is expressed as mean ± SEM (n = 3, **^#^***p* < 0.05 vs. the control group; **\****p* < 0.05 vs. the LPS group). Quercetin and luteolin could directly inhibit the IL-17 signaling pathway and simultaneously inactivate PI3K, inhibiting Akt and NF-κB, thereby preventing lung inflammation and treating CVA. **(B)** The dose-response curve of quercetin and luteolin on the growth of LPS-induced RAW264.7 cells. **(C)** The dose response curve of quercetin and luteolin on the growth of LPS-induced RAW264.7 cells.

Finally, we propose a novel MOA of Suhuang for treating CVA (Fig. 8C). We found that PI3K-Akt, NF-κB, and IL-17 signaling pathways were involved in the pathophysiological process of CVA. On the one hand, Phosphoinositide 3-kinases (PI3Ks) are involved in the immune response. The lipid products of PI3K activate Akt and then release NF-κB from the cytoplasm by phosphorylating a variety of TFs such as RELA and BCL3 ^52–54^. Activated NF-κB can induce many proinflammatory cytokines and produce lung inflammation. On the other hand, upon exposome stimulations, bronchial epithelial cells release IL-25, IL-33, and thymic stromal lymphopoietin. These cytokines promote the proliferation of Th17 cells and drive them to release IL-17, leading to lung inflammation ^55,56^. Therefore, we were able to predict the MOAs of the primary active compounds in Suhuang during the treatment of CVA: quercetin and luteolin could directly inhibit the IL- 17 signaling pathway and simultaneously inactivate PI3K, inhibiting Akt and NF-κB, and thereby preventing the occurrence of lung inflammation and treating CVA. This multi-compound, multi-target, and multi-pathway treatment of CVA provided a scientific explanation for the drug combination of Suhuang.

### 3.7 Association between TCM properties and the proximity distance of their targets with DEGs

The target-DEG shortest path of different targets varied from 0 to 1.287 (Fig. S4A), indicating that the interactions of these targets with DEGs differed. As the targets were predicted to interact with different ingredients from the nine herbs in Suhuang, we investigated whether the shortest path of target-DEG from different components was related to their respective herbs and properties (Fig. 9A).

**Fig. 9.**
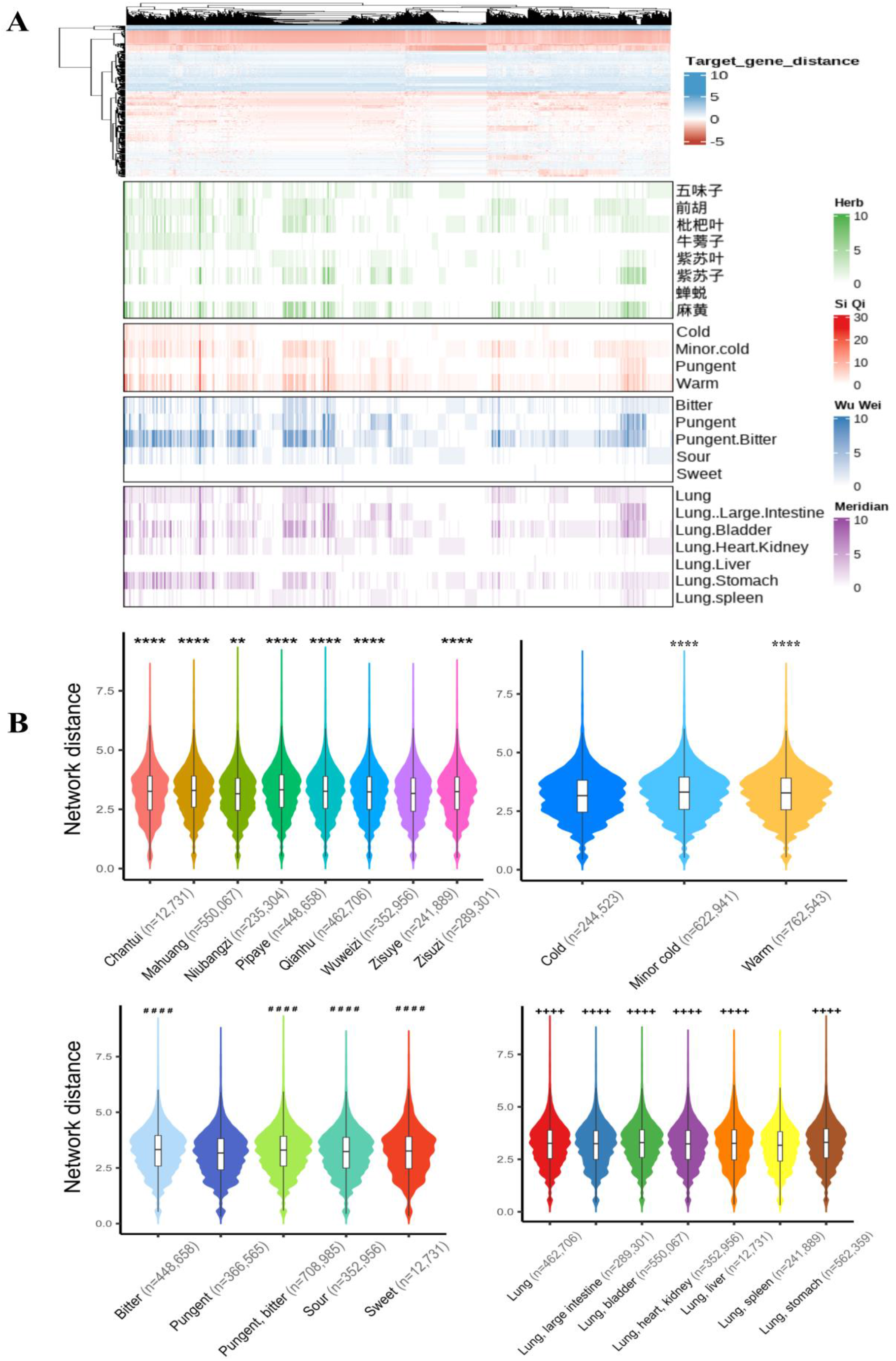
The calculated network distance of the DEGs of Suhuang and the targets from Suhuang and the TCM property distribution of targets. (A) The property distribution of targets in the target-DEG interactions. **(B)** The herb, nature, flavor, and meridian distributions of the targets in all the target-DEG interactions. *P-values* were calculated by <u>t.test.</u> \*\**p* < 0.01, \*\*\*\**p* < 0.0001 vs. the ZISUYE herb; \*\*\*\**p* < 0.0001 vs. the Cold nature; ^####^*p* < 0.0001 vs. the Pungent flavor; ^++++^*p* < 0.0001 vs. the Lung, spleen meridian.

Generally speaking, the target-DEG shortest path of targets from Zisuye was significantly lower than that of targets from other herbs (Fig. 9B). Additionally, Zisuye also tended to have more significant target-DEG shortest path in the transformed proximity network than other herbs, according to Fisher’s exact test (*p-value* = 3.56 × 10^-64^) (Fig. 10A). These results revealed the essential roles of targets from Zisuye, which were consistent with our prior knowledge that Zisuye was the monarch (“JUN”) medicine of Suhuang based on TCM theory. In terms of TCM properties, we found that the targets of cold nature, the pungent flavor, and the lung and spleen meridian tended to have significantly lower proximity with DEGs (*p-value* < 10^-4^) as well as more significant target-DEG pairs (Fisher’s exact test, *p-* value< 10^-4^) when compared to their counterparts in TCM theory (Fig. 10A).

**Fig. 10.**
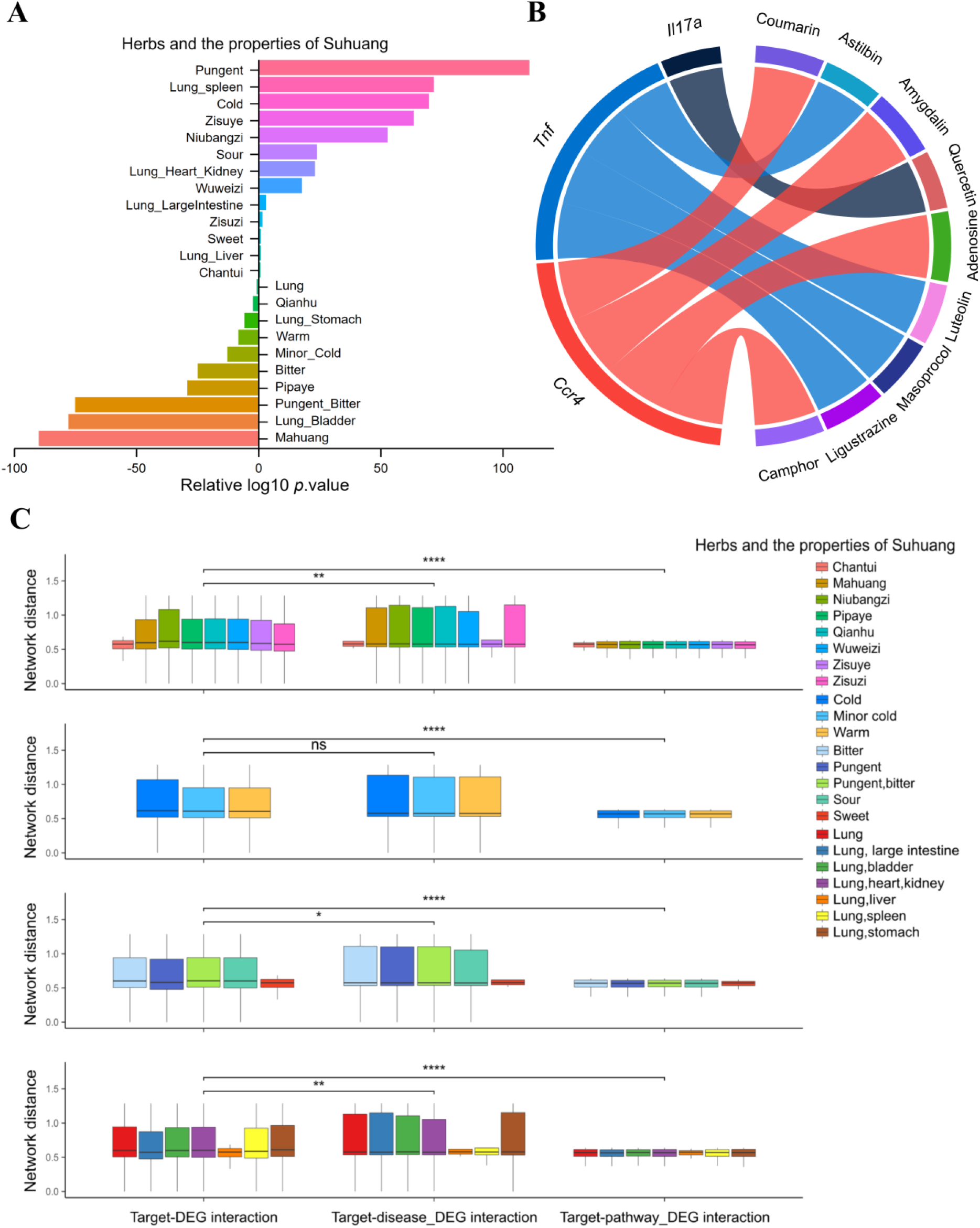
The important score of ingredients and target from LR methods. (A) The fisher extract test of the correlation between the target-DEG interactions with the significantly low network distance and the properties of targets. Relative log10 *p*.value > 0 & *p*.value < 0.05 suggested a positive correlation. **(B)** The herb, nature, flavor, and meridian distributions of the targets in the target-DEG, target-CVA DEG, and target-pathway DEG interactions with the significantly low network distance. *P-values* were calculated by <u>t.test.</u> \**p* < 0.05, \*\**p* < 0.01, \*\*\*\**p* < 0.0001. **(C)** The chord diagram of the active ingredients interacted with three CVA-related targets of Suhuang. Active ingredients (right) were connected to their targets (left) according to the compound-target interactions in Suhuang.

We found that the proximity of target-CVA gene and target-interesting pathway gene interactions was significantly lower than that of target-DEG interactions, suggesting that the significant targets we kept for network perturbation interacted strongly with CVA genes and were more likely to be the therapeutic targets (Fig. 10B). In particular, we identified six exciting targets which were not only DEGs of Suhuang but also related to CVA, including *Il17a*, *Tnf, Ifng*, *Il10*, *Ccr4*, and *Ido1*. Thus, we believe that Suhuang’s ingredients predicted to interact with these six targets deserve further verification (Fig. 10C).

More importantly, we found that targets from the lung and large intestine meridian exhibited significantly lower proximity in the transformed proximity network simultaneously (Fig. 10C). This reminded us that the lung and large intestine meridian played essential roles in Suhuang. This was consistent with the TCM theory of the “lung large intestine connection” ^57^. Specifically, based on our previous work, Suhuang could alleviate airway inflammation-associated lung injury by promoting Hepatocyte growth factor (HGF) secretion in an intestinal endocrine-dependent pathway. Moreover, the critical role of the lung meridian was in line with the tissue distribution of DEGs in the lung (Figure 4B).

In summary, by comparing the proximity of target-DEG interactions from the different herbs and properties of Suhuang, we found that the properties of Suhuang based on TCM theory could explain its mechanisms of treating CVA to some extent. We also reflected on the multi-component, multi-target, and multi-pathway characteristics of herbal medicine.

## 4. Discussion

Identifying the bioactive ingredients and critical targets of herbal medicine is difficult due to the lack of target information. Thanks to the development of computational biology and multi-omic methods, these large-scale data at different levels allow us to explore the scientific connotation of herbal medicine for treating diseases. To understand the molecular MOAs of herbal medicine treatments at the phenotype level, we proposed a novel method, Herb-CMap, to infer the therapeutic targets of herbal medicine using a gene perturbation network and the random walk algorithm. After finding potential targets, we worked backward to identify the bioactive ingredients according to the predicted ingredient-target pairs.

Herb-CMap completed the multiple data fusions required to uncover the MOAs of herbal medicine, such as PPI, gene profiling, disease genes associations, and herb-ingredient-targets. More importantly, advanced network proximity and network perturbation were used to integrate all the data. Our results showed that our network-based inference method could help to capture some bioactive targets that were not observed as significantly changed genes at the transcript level. For example, *Pik3cb*, a non-differentially expressed gene with a high rank by Diffusion, was closely interactive with adjacent target genes and showed a strong affinity with quercetin in Suhuang. Moreover, network proximity methods could help uncover the underlying association by considering the global topological characteristics. By calculating the network proximity of CVA-DEG, we quantified the importance of CVA-related genes in the gene perturbation of Suhuang. By calculating the target-DEG distance, we quantified the importance of targets of Suhuang for gene perturbation. However, although the “shortest step” network distance was applied in the previous model, the nodes and link attributions from the PPI network were not considered. To further optimize the algorithm, it is necessary to integrate the diversity of biological networks.

Herb-CMap can accurately predict targets and active ingredients, further accelerating the MOA exploration of herbal medicine and drug discovery from herbal medicine. In this case study, our method accurately predicted the known vital targets and the active ingredients of Suhuang and uncovered one novel ingredient. We successfully identified 17 active ingredients acting on the IL-17 signaling pathway and 21 working on the PI3K-Akt-NFκB signaling pathway, such as decursin and rosmarinic acid. It was reported that quercetin and luteolin could treat allergic asthma by acting as critical targets related to PI3K-Akt and IL-17 signaling pathways, which was also verified in vivo ^58,59^. Decursin could improve inflammatory diseases by regulating multiple pathways, including the PI3K-Akt- NFκB signaling pathway ^60^. Rosmarinic acid could also alleviate airway inflammation by inhibiting NFκB activation ^61^. Currently, few studies are available about the therapeutic effects of these compounds on asthma by regulating the IL- 17 signaling pathway. In addition, Herb-CMap also helped to discover a novel MOA: quercetin and luteolin could directly inhibit the IL-17 signaling pathway and simultaneously inactivate PI3K, inhibiting Akt and NF-κB, thereby preventing the occurrence of lung inflammation and treating CVA. In summary, Herb-CMap provides a novel insight into the deep understanding of how the complex systems of TCM prescriptions treat diseases by the MOAs of multi-compound, multi-target, and multi-pathway.

We also proposed a novel paradigm for deciphering TCM theories by investigating the relationships between our prioritized targets and the distributions of these properties. TCM theory was essential in using TCM formulae, such as nature, flavors, and meridians in TCM properties. First, for meridian theory, we found that the tissue expression attendance of DEGs followed the lung meridian of Suhuang. Second, Zisuye had closer interactions with DEGs than other herbs that are considered to play significant therapeutic effects, which is in line with its monarch role in TCM theory. Studies have shown that the extraction of Zisuye could effectively alleviate airway inflammation and high airway responsiveness in the asthma mouse model induced by ovalbumin ^62,63^. Specifically, Zisuye contains various active ingredients, including flavonoids and phenolic acids such as luteolin and rosmarinic acid, with anti-inflammatory activity ^64,65^. These ingredients in Zisuye may be the therapeutic material basis in Suhuang for treating CVA. Third, in terms of the “JUN-CHEN-ZUO-SHI” theory of combination, we found that NIUBANGZI and WUWEIZI had significant target-DEG shortest path according to Fisher’s exact test, with *p* values 1.63 × 10^-53^ and 2.08 × 10^-18^, respectively. Accordingly, this also suggested that the active compounds corresponding to the crucial targets from NIUBANGZI and WUWEIZI had therapeutic roles in Suhuang for treating CVA, which was also consistent with our prior knowledge that NIUBANGZI and WUWEIZI were the minister (“CHEN”) medicines based on TCM theory. Additionally, the main active components of NIUBANGZI and WUWEIZI, such as arctigenin and schizandrin, were found to relieve airway inflammation and treat asthma ^66,67^. Fourth, the results of the property distributions of Suhuang’s targets based on TCM theory were also consistent with modern scientific research. If the transformed proximity network was noticed, the proximity of targets of the pungent flavor was also lower than that of other flavors (Fig. 9B). It had previously been reported that transient receptor potential (TRP) channels were responsible for various sensory reactions, including heat, cold, pain, stress, vision, and taste ^68^.

For example, TRP channel subfamily V-1 (TRPV1) was sensitive to capsaicin interrelated with the pungent flavor, and TRP channel subfamily M-8 (TRPM8) was related to the cold sensory ^69–71^. They also played a role in respiratory diseases, nervous and mental diseases, diabetes, and cancer. Therefore, the property classifications of Suhuang might be associated with regulating these proteins. To sum up, our Herb-CMap could help us better understand TCM theory in formulae at the molecular level^72^.

There are some limitations in this study. For example, our network perturbation method considered first-level neighborhood genes in PPI, namely LR. Many other advanced network perturbation methods are worth trying.

In summary, this study addresses the challenge of identifying bioactive ingredients and key targets in herbal medicine due to limited target information. Leveraging computational biology and multi-omic methods, we introduce a novel approach called Herb-CMap to understand the molecular mechanisms of herbal medicine treatments. Herb-CMap integrates various data sources, including protein-protein interactions, gene profiling, disease-gene associations, and herb-ingredient- targets, employing network proximity and perturbation methods to uncover therapeutic targets not detected at the transcript level. This approach accurately predicts targets and active ingredients, facilitating the exploration of herbal medicine mechanisms and drug discovery, exemplified by identifying active ingredients in Suhuang for treating Cough Variant Asthma and offering insights into TCM theory.

## Acknowledgments

This work was supported by the “Double First-Class” University Project (CPU2018GF05) from China Pharmaceutical University and the Program for Innovative Research Team of Jiangsu Province.

---

**Fig. S1.**
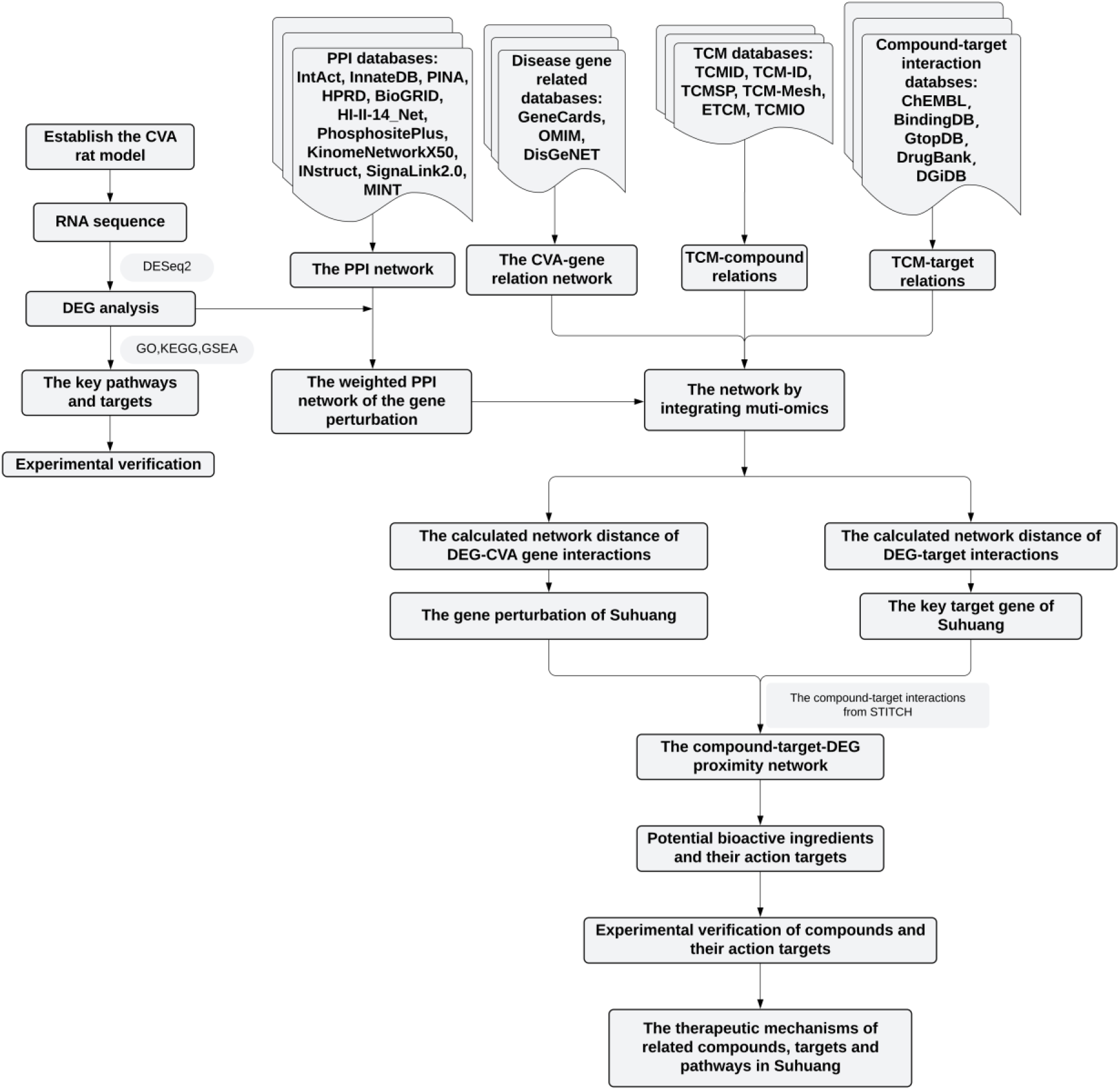
The workflow of Herb-CMap.

**Fig. S2.**
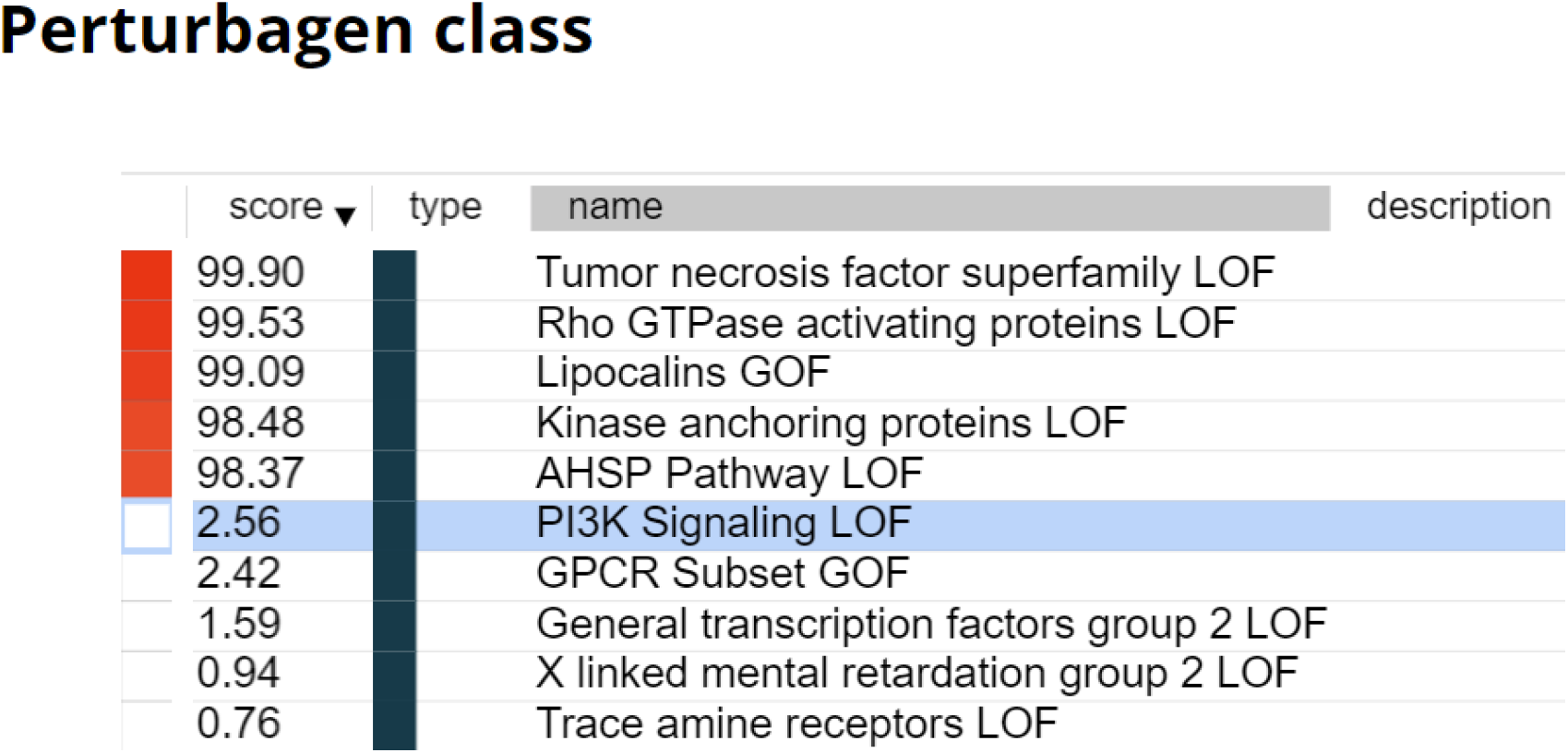
The connection map of DEGs for MOAs.

**Fig. S3.**
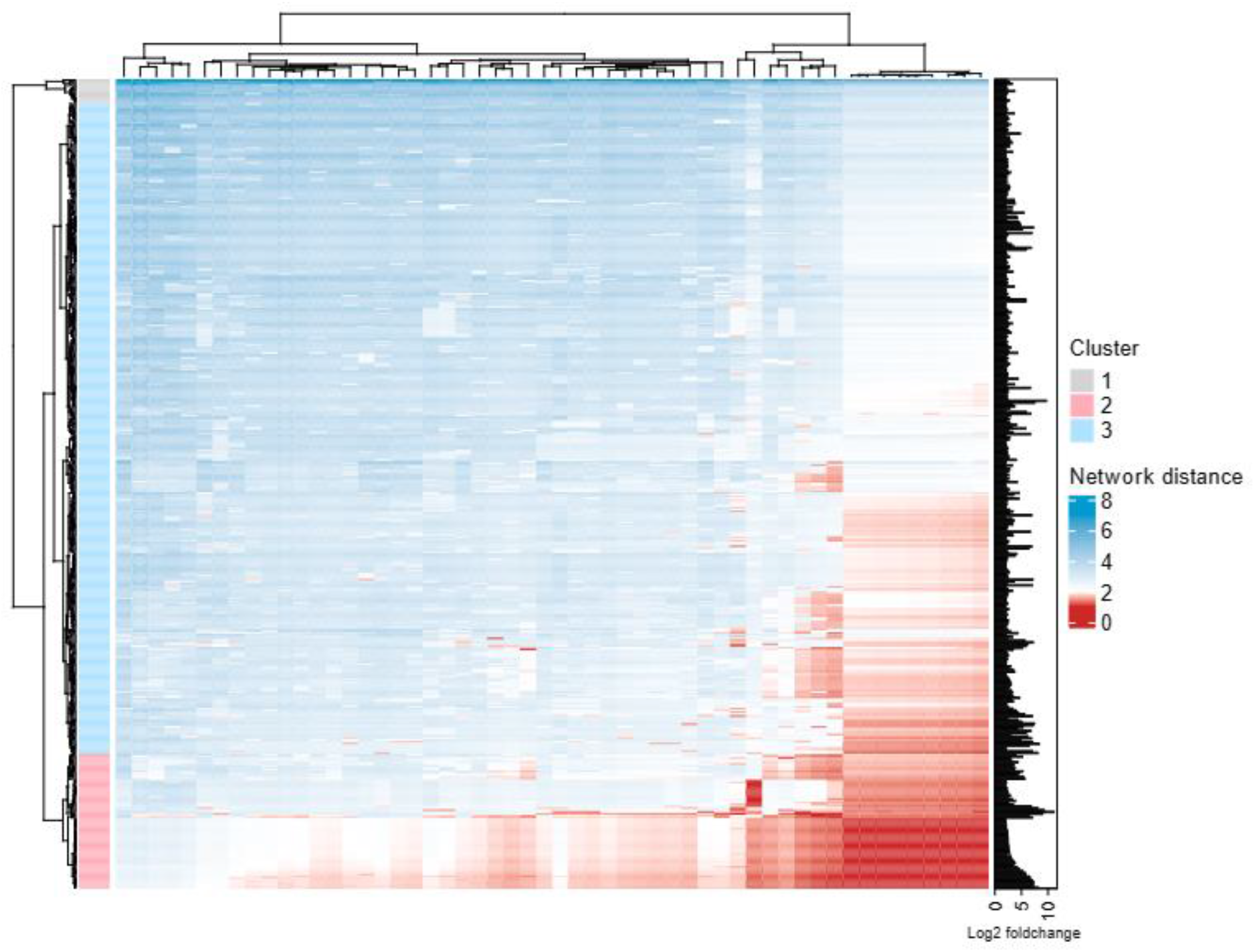
Based on the network proximity model, The calculated network distance between the DEGs of Suhuang and the CVA-related genes.

**Fig. S4.**
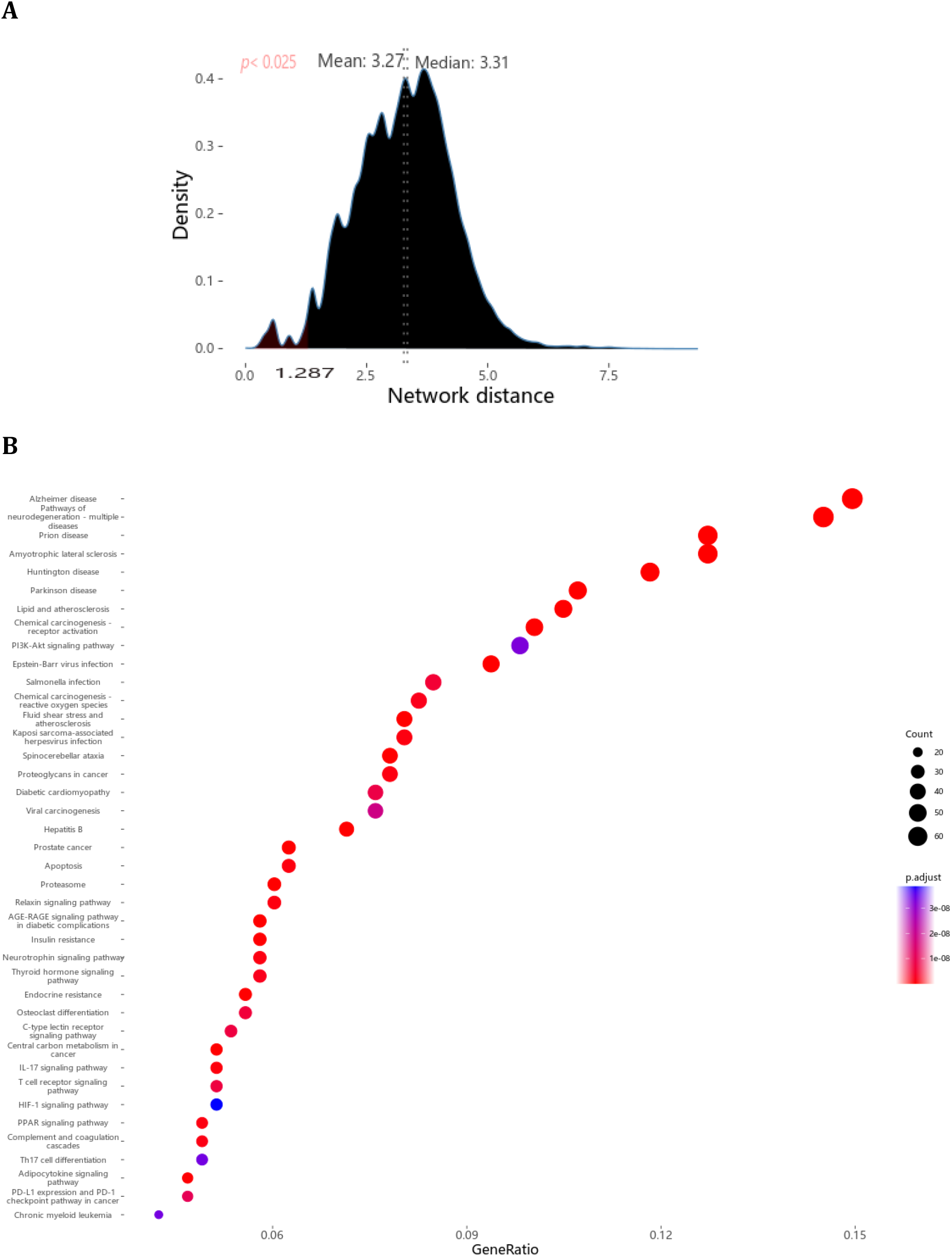
The target-DEGs distance analysis. (A) The network distance distribution of all the target-DEG interactions (n = 928,485). The red part of the density plot was the target-DEG interactions with the significantly low network distance (n = 19,009) based on the one-sided test of normal distribution (*z* < 0, *p* < 0.025). **(B)** The topmost significant signaling pathways were enriched by introducing 499 targets that were not DEGs into the KEGG pathway enrichment analysis.

## Notes

### Competing Interest Statement

The authors have declared no competing interest.

### Summary of Updates

small change for manuscript yo make the manuscript more readable

